# Characterizing the extracellular matrix transcriptome of endometriosis

**DOI:** 10.1101/2023.03.08.531805

**Authors:** Carson J. Cook, Kaitlin C. Fogg

## Abstract

In recent years, the matrisome, a set of proteins that make up the extracellular matrix (ECM) or are closely involved in ECM behavior, has been shown to have great importance for characterizing and understanding disease pathogenesis and progression. The matrisome is especially critical for examining diseases characterized by extensive tissue remodeling. Endometriosis is characterized by the extrauterine growth of endometrial tissue, making it an ideal condition to study through the lens of matrisome gene expression. While large gene expression datasets have become more available, and gene dysregulation in endometriosis has been the target of several studies, the gene expression profile of the matrisome specifically in endometriosis has not been well characterized. In our study, we explored three Gene Expression Omnibus (GEO) DNA microarray datasets containing endometriosis and healthy samples of eutopic endometrium. We established that matrisome gene expression alone can stratify healthy and endometriosis samples and identified the matrisome genes and gene networks that hold inferential significance for the onset and progression of endometriosis. Furthermore, we found that menstrual cycle phase accounted for over a third of the variance of matrisome gene expression within the samples. Taken together, these findings may aid in developing future *in vitro* models of disease and identifying novel treatment strategies for this underserved patient population.

## Introduction

Endometriosis affects approximately 10-15% of women of reproductive age and is characterized by the growth of ectopic endometrium.^1–4^ This can cause women to suffer from chronic, sometimes debilitating, pain, infertility, and other dysfunction of reproductive organs.^1, 3, 5^ While the underlying cause of endometriosis remains unknown, tissue remodeling is critical to the pathogenesis and progression of this disease.^6, 7^ Tissue remodeling is a complex and dynamic process, involving both extracellular matrix (ECM) deposition as well as ECM degradation.^8^ While individual components of the ECM and ECM-affiliated cytokines have been investigated, the ECM and ECM affiliated proteins form a complex interconnected network of over 1,000 genes known as the matrisome.^9^ Thus, a holistic yet targeted evaluation of the entire endometriosis matrisome holds the potential to elucidate specific microenvironmental cues involved in the underlying pathogenesis as well as perpetuation of endometriosis.

Though endometriosis has been shown to have a strong association with heredity and family clustering, it is not hereditary in a predictable Mendelian manner.^2, 4^ Transcriptomics analyses, which quantify and assess gene expression in disease and healthy tissue, are well-suited for characterizing gene expression in endometriosis pathophysiology. However, only one large-scale transcriptomic analysis has been performed using DNA microarrays to study endometriosis,^4^ assessing global gene expression and focusing on immune infiltration. To our knowledge, no existing studies have performed a targeted analysis of matrisome gene expression in endometriosis using large gene expression datasets of endometriosis tissue samples. Thus, the goal of this study was to establish the significance of the matrisome in characterizing endometriosis and identify key matrisome components which have inferential value with respect to the onset and progression of endometriosis.

In this study, we unified publicly available whole transcriptome microarrays from normal and endometriosis samples of eutopic endometrium. We employed a variety of statistical and machine learning methods to explore dysregulation of genes in endometriosis and identify the matrisome genes, gene networks, gene ontology (GO) terms,^10^ and pathways^11^ which have significance for the onset and progression of endometriosis. We found that matrisome gene expression effectively stratified endometriosis and normal tissue and that ECM-related GO terms were highly enriched among differentially expressed matrisome genes. Additionally, we found that menstrual cycle phase accounted for over a third of the matrisome gene expression variance, thus we needed to separate the data by menstrual cycle phase before performing differential expression analysis, statistical and machine learning modeling, and enrichment analysis. From these approaches we identified matrisome genes and gene networks with inferential significance for endometriosis stage.

## Materials and Methods

### Data sources and preprocessing

All data preprocessing was done using the R programming language.^12^

### Gene Expression Omnibus data

The Gene Expression Omnibus (GEO) database (http://ncbi.nlm.nih.gov/geo/) was accessed to retrieve four datasets using the same search criteria, and subsequent filtration methods, as Poli-Neto *et al*.^4^ The four datasets which were retrieved were from GSE4888,^13^ GSE6364,^14^ GSE7305,^15^ and GSE51981.^16^ However, the data corresponding to GSE7305 were excluded because the samples were paired (disease and normal tissue samples taken from the same patient). It was also found that the tissue samples were collected from the ovaries rather than the uterus, whereas the other studies included only uterine samples. Each of these datasets was made up of samples assessed using the Affymetrix Human Genome U133 Plus 2.0 Array (HG-U133 Plus 2, Affymetrix, Santa Clara, CA).^4^ The data were then loaded and normalized via the robust multiarray average method (RMA) using the *Affy* package.^17–19^ Finally, the data were batch corrected using the *comBat* function from the *sva* package,^20^ using GSE4888 as the reference batch. This was the approach that yielded the best empirical results in terms of removing batch effects (along with batch interactions) and the most variance attributable to disease condition and menstrual cycle phase. The weighted proportion of variance for the effects factors of interest was computed using principal variant component analysis (PVCA), which combines principal component analysis (PCA) and variance component analysis (VCA) to determine the amount of variance in the data attributable to specified variables.^21^ The *pvcaBatchAssess* function from the *pvca* package was used with the parameter *threshold* assigned a value of 0.6. This function allowed us to assess the relative amount of variance, divided among factors of interest using principal components, which explained 60% of the variance in the data.^22^ Clinical data were retrieved by downloading the series matrix files, loading them using the *getGEO* function from the *GEOquery* package,^23^ and then performing necessary data cleaning. The endometriosis samples in the data were classified as min/mild or moderate/severe, but “min/mild” is referred to as simply “mild” throughout this study.

### The matrisome database

The human matrisome database, compiled by Naba *et al*., was retrieved from their online repository (http://matrisomeproject.mit.edu/other-resources/human-matrisome) on 2020/07/21.^24^ Genes classified as “retired” were filtered out, yielding a master list of 1027 genes, 964 of which were present in the GEO datasets. Of the genes in the combined dataset, the divisions consisted of “Core matrisome” (*n* = 258) and “Matrisome-associated” (*n* = 706). Matrisome categories included: collagens, ECM glycoproteins, ECM regulators, ECM-affiliated proteins, proteoglycans, and secreted factors (**Table 1**).

**Table 1.** Matrisome category counts in master list and in dataset. Counts of matrisome genes in each matrisome category in the matrisome master list and in our dataset. Of the 1,027 non-retired matrisome genes in the human matrisome master list, 964 were tested for using the Affymetrix Human Genome U133 Plus 2.0 Array chip.

| Division | Category | Master list count | Dataset count |
| --- | --- | --- | --- |
| Core matrisome | Collagens | 44 | 44 |
| Core matrisome | ECM glycoproteins | 195 | 179 |
| Core matrisome | Proteoglycans | 35 | 35 |
| Matrisome-associated | ECM regulators | 238 | 230 |
| Matrisome-associated | ECM-affiliated proteins | 171 | 151 |
| Matrisome-associated | Secreted factors | 344 | 325 |

### Dimensionality reduction

Principal component analysis (PCA) was performed using the *prcomp* function from the *stats* package in *R*.^12^ PCA was used to explore batch and biological effects in the data.

### Stratification of endometriosis and normal tissue

Within each phase, elastic net logistic regression models were trained on the RMA matrisome data.^25^ Elastic net regression utilizes the objective function

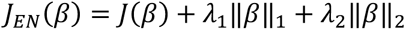

where *J*(*β*) is a less complex loss function, and the parameters *λ*_1_ and *λ*_$_ control the proportion of L^1^ (lasso) and L^2^ (ridge) regression penalization to use. Within each phase, these models were trained to classify samples as endometriosis as normal or endometriosis tissue. Model performance was measured using balanced classification accuracy.^26^ This scoring method utilizes observation weights defined according to

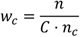

where *w_c_* is the weight assigned to observations from class *c*, *n* is the total number of observations, *C* is the total number of classes (factor levels of the response), and *n_c_* is the number of observations in class *c*.^27^ Observation weights sum to 1, ensuring that models are penalized equally for bad performance in any class regardless of the imbalanced representation of classes. Using sequential model-based optimization,^28^ these models were optimized based on 5-fold cross-validation scores. All features (genes) were standardized to have a mean of 0 and a standard deviation of 1. The *scikit-learn* implementation of logistic regression was used,^27^ and sequential model-based optimization was performed using *gp_minimize* from *scikit-optimize*.^29^ These packages are open source and available for the Python programming language, which was used for this portion of the study.

### Differential gene expression analysis

Differential gene expression (DGE) analysis was conducted on the full set of RMA normalized gene expression counts using the *limma* package in *R*.^30^ All genes with expression units larger than *log*_$_(50) (*Affy*’s *rma* function produces results in *log*_$_) in at least 25% of samples were considered sufficiently expressed. A permissive expression cutoff was chosen so that matrisome genes were filtered at similar rates in each category to genes overall (**Table 2**).

**Table 2.** Gene filtration rates during differential gene expression analysis. Number of genes overall and number of matrisome genes within each matrisome category before and after expression level filtration as part of DGE analysis pipeline.

| **All genes (n = 21,407)** |  |  |  |  |
| --- | --- | --- | --- | --- |
|  |  | **Dataset count** | **Post-filtration count** | **Retention rate** |
| All genes |  | 21,407 | 17,168 | 80% |
| **Matrisome genes (n = 964)** |  |  |  |  |
| **Division** | **Category** | **Dataset count** | **Post-filtration count** | **Retention rate** |
| Core matrisome | Collagens | 44 | 36 | 82% |
| Core matrisome | ECM glycoproteins | 179 | 149 | 83% |
| Core matrisome | Proteoglycans | 35 | 27 | 77% |
| Matrisome-associated | ECM regulators | 230 | 189 | 82% |
| Matrisome-associated | ECM-affiliated proteins | 151 | 124 | 82% |
| Matrisome-associated | Secreted factors | 325 | 235 | 72% |
| Combined |  | 964 | 760 | 79% |

DGE analysis was then performed by phase, and genes with a log fold-change of (1.5) and adjusted *p*-value less than 0.05 were considered differentially expressed. Adjusted *p*-values were computed using the Benjamini-Hochberg false discovery adjustment method.

### Univariable and multivariable statistical analyses

Several univariate and multivariable statistical analyses were performed on the RNA normalized gene expression data (endometriosis samples only). The stage-wise DGE analysis was performed using all genes to allow for better estimation of global parameters. The other analyses were performed on the matrisome genes alone. False discovery rates were estimated for statistical tests using computed *q*-values.^31^ Similar to the endometriosis versus normal tissue DGE analysis, the data were stratified by phase for each analysis, yielding separate results for each menstrual cycle phase.

### Stage-wise differential gene expression analysis

Differential gene expression analysis was performed between endometriosis samples from the two different endometriosis stages in the dataset, “minimal/mild” and “moderate/severe.” The same minimum expression threshold as the endometriosis versus normal DGE analysis was used. This analysis was performed on the set of all genes, at which point the results were filtered to include only matrisome genes. Differential expression was determined using the same adjusted *p*-value and log fold-change cutoffs as the endometriosis versus normal DGE analysis. The *limma* package was used for this analysis.

### Point-biserial correlation

Point-biserial correlation is mathematically equivalent to the Pearson correlation between a continuous and dichotomous variable.^32^ Endometriosis stage was coded as an indicator variable, with a value of 1 corresponding moderate/severe endometriosis and 0 corresponding to “minimal/mild” endometriosis. Genes were deemed significant if their Student asymptotic *q*-values were below the significance threshold (*q* < 0.05).^33^ Student asymptotic *p*-values were computed using the *corPvalueStudent* function from the *WGCNA* package and then adjusted.

Endometriosis stage-predictive penalized logistic regression

Multivariable L^1^ penalized logistic regression models were fit to the endometriosis data in each phase. The models were fit using the *glmnet* package in *R*.^34^ The models were optimized for parsimony which is done by tuning the value of *λ* in the L^1^ penalty objective function

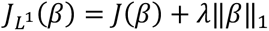

where *J*(*β*) is some simpler loss function (misclassification rate, for example) and ‖*β*‖_1_is the L^1^ norm of the model parameters. The models were fit using a 1-dimensional grid search over values of *λ*, and the least parsimonious model (smallest value of *λ*) which scored within 1 standard error of the best performing model was selected for each phase. This was done to avoid using models which selected too few matrisome genes, since these results were then filtered to include only DEMGs. The models were fit using 5-fold cross-validation using the *cv.glmnet* function from the *glmnet* package with arguments *family* set to “binomial” and *type.measure* set to “class.” Balanced class-weighting (as in the endometriosis/normal stratification models) was used to ensure models did not favor performance in only the majority class.

### Weighted gene correlation network analysis

Univariable and multivariable analyses assessed a gene’s independent association with endometriosis stage or a gene’s ability to serve as a proxy for a set of other co-expressed genes. In contrast, weighted gene correlation network analysis (WGCNA) identified genes which, as a cluster, demonstratd a significant link to endometriosis progression.^33^ This was achieved by establishing unsigned gene co-expression modules, where co-expression is estimated using unsigned topological overlap measures.^35^ Each module was then represented using the module’s eigengene, which was the first principal component representation of the gene expression of all genes assigned to that module.^33^ These eigengenes were then be correlated with endometriosis stage. Finally, a module’s constituent genes were identified as significant based on correlation tests with their respective eigengenes.

WGCNA was performed according to the instructions provided by the package authors.^36^ First, the data were stratified by phase. Then a topological overlap measure matrix was constructed over the matrisome gene expression values using a minimum soft power threshold which yielded a scale-free topological overlap metric (TOM) greater than 0.8,^35^ representing a gene-wise estimate of unsigned co-expression among matrisome genes. Hierarchical clustering was then performed on this matrix, and modules with a correlative distance of 0.25 or less (i.e., module correlation of 0.75 or more) merged. The modules found using this method were then related to endometriosis severity via point-biserial correlation. Matrisome genes that belonged to a module which was significantly correlated with endometriosis stage (Student asymptotic *q* < 0.05) and showed significant correlation with their respective module eigengenes (Student asymptotic *p* < 0.05) were deemed significant. All WGCNA was performed using *R* code and functions from the *WGCNA* package in *R*.^33^

### Enrichment analysis

Gene set and pathway enrichment analysis was performed using the *clusterProfiler* package in *R*.^37^ For gene set enrichment analysis, the function *enrichGO* was used to find enriched gene ontologies (GO) among significant genes.^38^ For pathway enrichment analysis, the function *enrichKEGG* was used to find KEGG pathways that were enriched among significant genes.^11^ Gene function and pathway significance was determined based on the *q*-value reported in the results of each function (*q* < 0.05).

## Results

### Sources of variance

To examine the sources of variance in our dataset, we used principal component analysis (PCA). Because the data we used came from three different clinical studies, we assessed the level of variance attributable to technical differences in the data collection processes. To correct for these sources of technical variance, often called “batch effects,” we performed batch correction using an empirical Bayes method (**Figure 1**).^39^ The data used in our study consisted of tissue samples from the proliferative, early-secretory, and mid-secretory phases of the menstrual cycle from patients with and without a clinical diagnosis of endometriosis.

**Figure 1.**
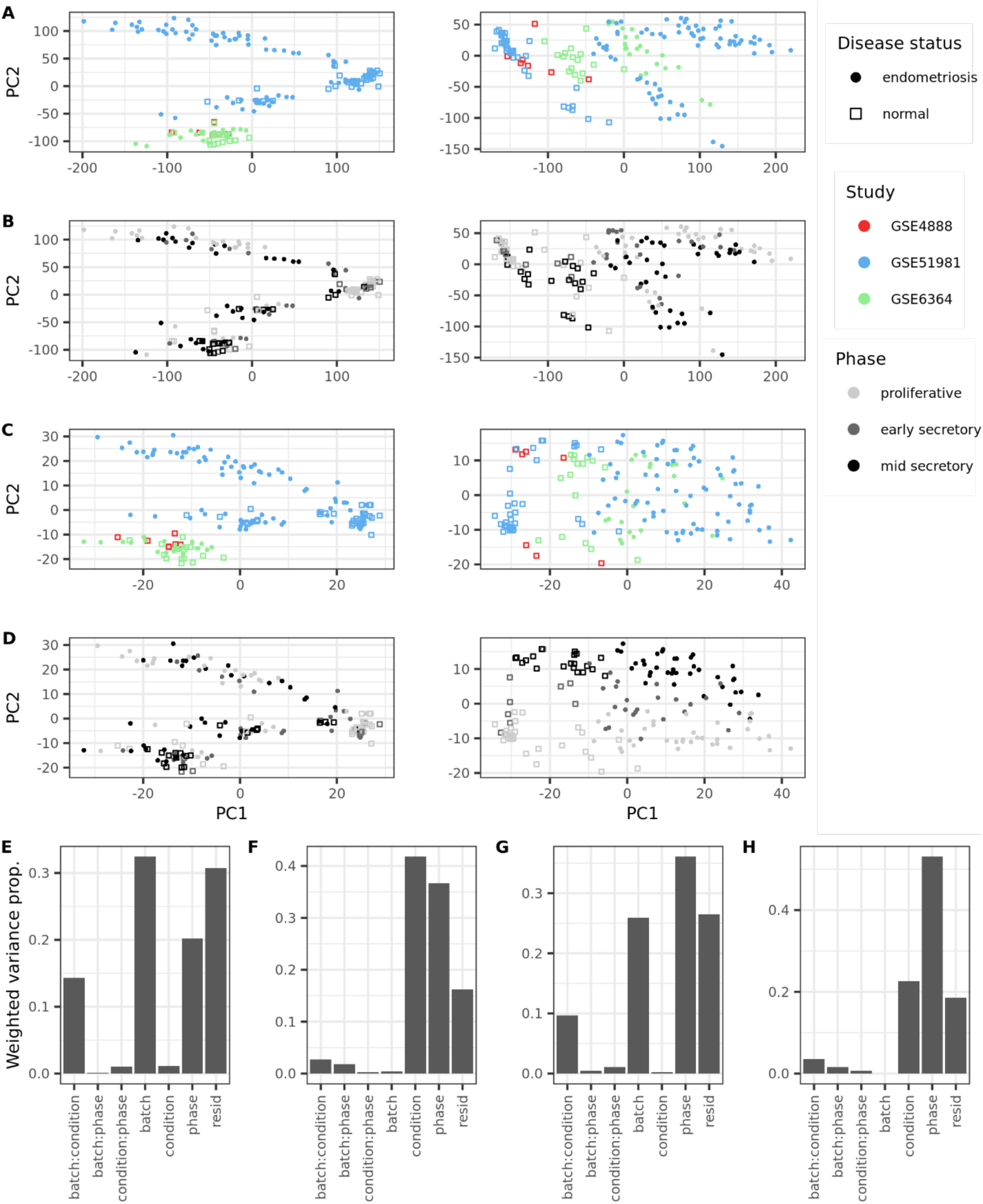
Sources of variance within the data. **(A)** Principal component analysis (PCA) of the full gene expression data (*n_genes_* = 21,407) before (left) and after (right) batch correction. Each clinical study is distinguished by color (GSE4888, red; GSE51981, blue; GSE6364, green). Percent variance explained: left (PC1, 33.6%; PC2, 24.3%), right (PC1, 49.4%; PC2, 9.3%). **(B)** PCA of the full gene expression data before (left) and after (right) batch correction. Each menstrual cycle phase is distinguished by shade (proliferative, light gray; early-secretory, dark gray; mid-secretory, black). Percent variance explained: left (PC1, 33.6%; PC2, 24.3%), right (PC1, 49.4%; PC2, 9.3%). **(C)** PCA of the matrisome gene expression data (*n_mat. genes_* = 964) before (left) and after (right) batch correction. Each clinical study is distinguished by color. Percent variance explained: left (PC1, 27.5%; 20.6%), right (PC1, 41.6%; PC2, 10.0%). **(D)** PCA of the matrisome gene expression data before (left) and after (right) batch correction. Each menstrual cycle phase is distinguished by shade. Percent variance explained: left (PC1, 27.5%; 20.6%), right (PC1, 41.6%; PC2, 10.0%). **(E)** Principal variant component analysis (PVCA) of global gene expression before batch correction. For PVCA, the *x*-axis corresponds to the factors of interest and their linear interaction terms. **(F)** PVCA of global gene expression after batch correction. **(G)** PVCA of matrisome gene expression before batch correction, **(H)** PVCA of matrisome gene expression after batch correction. Sample sizes by study: GSE4888 (*n_normal_* = 7, *n_disease_* = 0), GSE51981 (*n_normal_* = 34, *n_disease_* = 75), GSE6364 (*n_normal_* = 16, *n_disease_* = 21). Sample sizes by phase: proliferative (*n_normal_* = 28, *n_disease_* = 35), early-secretory (*n_normal_* = 9, *n_disease_* = 24), mid-secretory (*n_normal_* = 20, *n_disease_* = 37).

We examined the variance in our data due to menstrual phase and disease status before and after batch correction. For the global gene expression data (*n_genes_* = 21,407), before batch correction the first principal component captured 34% of the variance and appeared to separate samples mostly by disease status, whereas the second component captured 24% of the variance and appeared to primarily separate samples by study. After batch correction, the first component explained 49% of the variance in the data and clearly separated samples by disease status. The second component explained only 9% of the variance and seemed to somewhat separate samples by menstrual cycle phase (**Figure 1A, B**). When we examined the gene expression of the matrisome alone (*n_mat.genes_* = 964), before batch correction the first principal component captured 28% of the variance and appeared to separate samples mostly by disease status, while the second component captured 21% of the variance and separated samples primarily by study. After batch correction, the first component accounted for 42% of the variance and separated samples by disease status, while the second component explained 10% of the variance and appeared to mostly separate samples by menstrual cycle phase (**Figure 1C, D**). Principal variant component analysis (PVCA) was then used to reduce the gene expression data to the first principal components which accounted for 60% of the variance, then evaluate relative proportions of variance within these principal components due to clinical study, disease status, menstrual phase, and their interactions. PVCA was performed before and after batch correction for global (**Figure 1E, F**) and matrisome (**Figure 1G, H**) gene expression. Before batch correction, batch (study) and batch interaction terms accounted for approximately 47% of the variance in the global gene expression data and 36% in the matrisome gene expression data. After batch correction, these percentages were reduced to approximately 5% in both the global and matrisome gene expression data. We found that disease status accounted for the most variation in the batch corrected global expression data (42%), closely followed by menstrual cycle phase (37%). For the batch corrected matrisome expression data, menstrual cycle phase accounted for the most variation (53%), followed by disease status (23%). The high level of variance attributed to menstrual cycle phase in the matrisome gene expression data was unsurprising, given that the endometrium is highly dynamic and heterogeneous between phases of the menstrual cycle ^13, 40^.

### Stratification by menstrual phase

The demonstration of extensive variance attributable to menstrual cycle phase prompted us to adopt a similar approach to Poli-Neto *et al.* and stratify the data by menstrual cycle phase before performing our differential expression analysis, statistical and machine learning modeling, and enrichment analysis ^4^. However, in each of these analyses, it became apparent that the differentially expressed and otherwise significant matrisome genes in the early and mid-secretory phases were almost entirely subsets of those found to be differentially expressed or significant in the proliferative phase (**Figure 2**).

**Figure 2.**
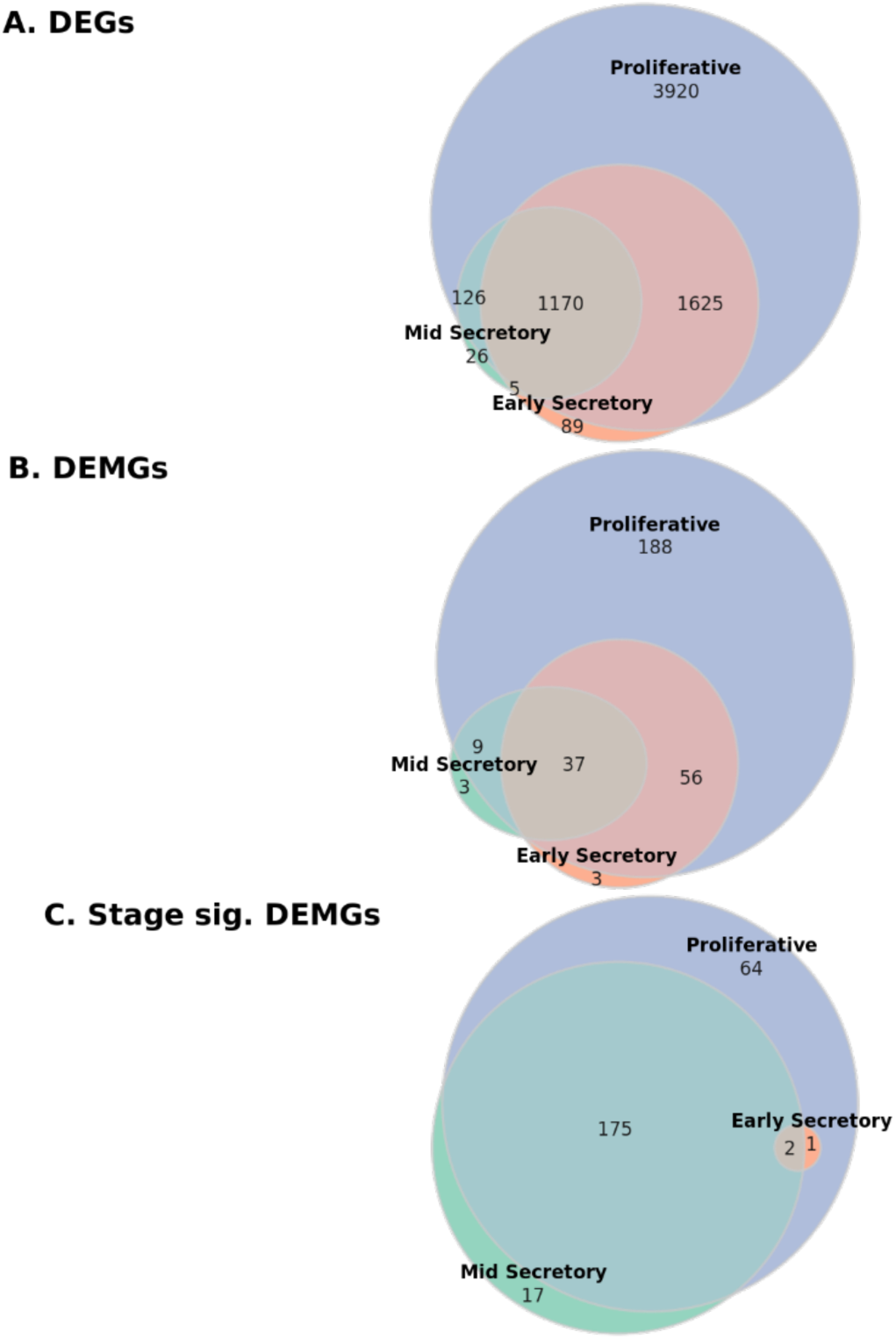
Overlaps of significant genes between phases. Inter-phase overlaps within the set of **(A)** genes which were differentially expressed between normal and endometriosis tissue (DEGs) (*n* = 6961), **(B)** matrisome genes which were differentially expressed between normal and endometriosis tissue (DEMGs) (*n* = 296), and **(C)** DEMGs which were significant with respect to endometriosis stage (*n* = 259).

Furthermore, when there was overlap between phases, the directionality of significant matrisome gene influence (e.g., fold-change sign) was also virtually identical between phases for all analyses. Only 1 of 6,961 unique DEGs was found to be differentially expressed in endometriosis in an inconsistent direction between phases, and only 1 of 259 DEMGs later found to be significantly related to endometriosis stage was directionally inconsistent between phases (**Table 3**). For this reason, we decided to conduct the various analyses separately within each phase, then perform a union operation on the results of the various analyses – pooling all unique genes significant in any phase – before conducting enrichment analyses.

**Table 3.** Mismatched gene significance between phases. Among genes shown to be differentially expressed between endometriosis and normal tissue, *ATP12A*, a non-matrisome gene, had differing significance between phases. Among differentially expressed matrisome genes (DEMGs) with significance for endometriosis stage, *ANXA4* showed a different form of significance in different phases. Sample sizes: CESC (*n_normal_* = 13, *n_tumor_* = 259), UCEC (*n_normal_* = 105, *n_tumor_* = 141), and UCS (*n_normal_* = 105, *n_tumor_* = 47).

|  | Phase |  |  |
| --- | --- | --- | --- |
| Gene ID | Proliferative | Early secretory | Mid Secretory |
| ***Endometriosis vs. normal differential gene expression analysis (all genes)*** |  |  |  |
| ATP12A | Upregulated | Not sig. | Downregulated |
| ***Stage-wise differential gene expression analysis (matrisome)*** |  |  |  |
| ANXA4 | Upregulated | Downregulated | Upregulated |

### Differentially expressed genes between endometriosis and normal uterine tissue samples

To investigate the importance of the matrisome in characterizing endometriosis, we performed differential gene expression (DGE) analysis on the full set of genes in the dataset (*n_genes_* = 21,407), comparing endometriosis samples (*n_proliferative_* = 35, *n_early-secretory_* = 24, *n_mid-secretory_* = 37) to normal tissue samples (*n_proliferative_* = 28, *n_early-secretory_* = 9, *n_mid-secretory_* = 20), examined differential expression rates among matrisome genes (*n_mat.genes_* = 964), and performed functional enrichment analysis on the full list of differentially expressed genes (DEGs) to observe whether ECM-related gene ontology (GO) terms were enriched. The DGE analysis was performed separately for samples in each phase, and 6,841, 2,889, and 1,327 differentially expressed genes (DEGs) were found in the proliferative, early-secretory, and mid-secretory phases, respectively (**Figure 3A, Table 4**). Next, we filtered these results to include only the matrisome genes and identified 290, 96, and 49 differentially expressed matrisome genes (DEMGs) in the proliferative, early-secretory, and mid-secretory phases, respectively (**Figure 3B, Table 4**).

**Figure 3.**
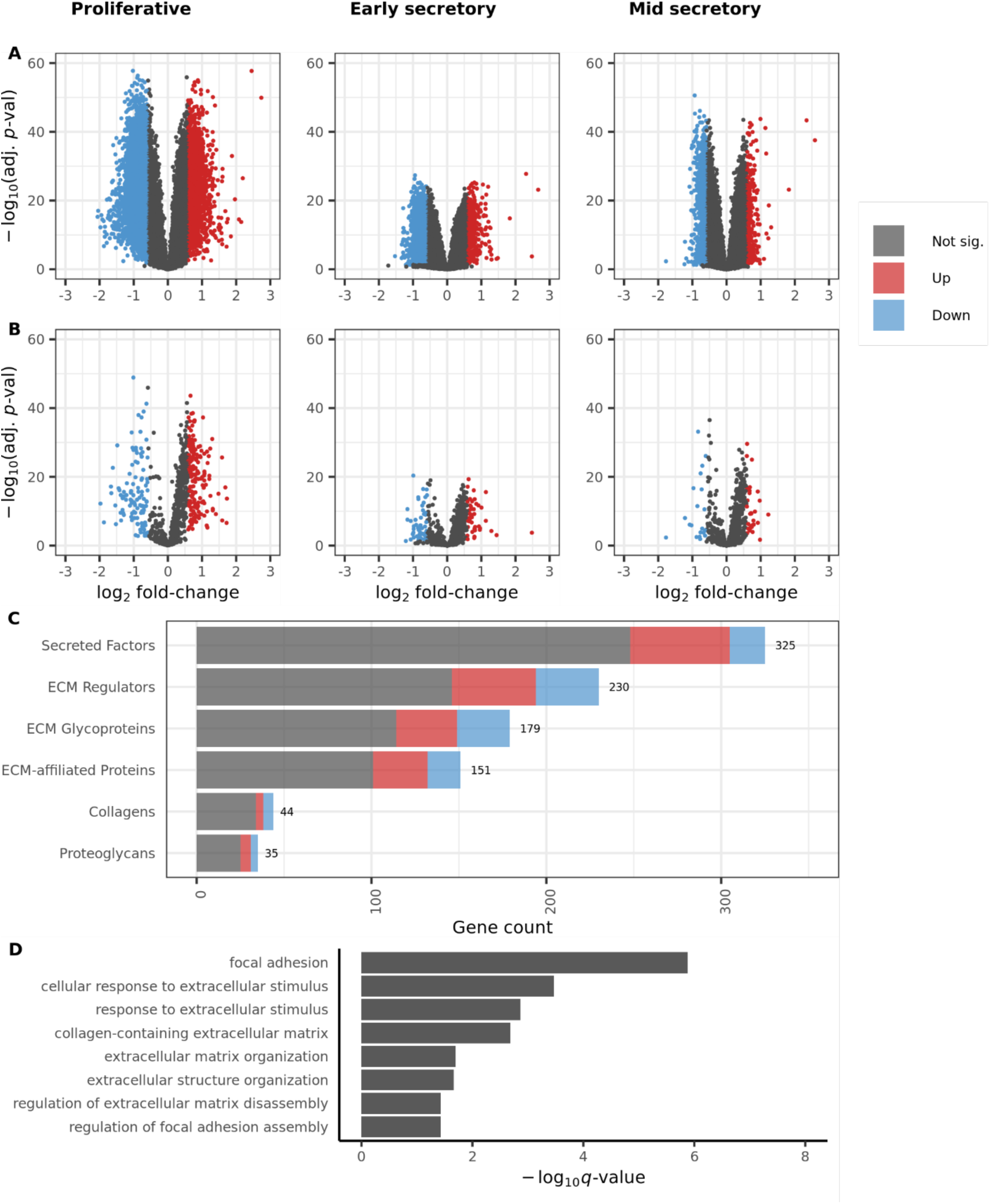
Differential gene expression within each phase and functional enrichment analysis of combined results. Differentially expressed genes in each phase among **(A)** all genes and **(B)** matrisome genes. **(C)** Breakdown of differentially expressed matrisome genes (union of results in all phases) by matrisome category. **(D)** Results of functional enrichment analysis for all genes, with respect to ECM-related gene functions. Gene counts in dataset: all genes (*n_genes_* = 21,415) and matrisome genes (*n_mat. genes_* = 964). Sample sizes by phase: proliferative (*n_normal_* = 28, *n_disease_* = 35),early-secretory (*n_normal_* = 9, *n_disease_* = 24), mid-secretory (*n_normal_* = 20, *n_disease_*= 37).

**Table 4.** Differentially expressed gene counts. Counts and percentages of differentially expressed (DE) genes among all genes (*n* = 21,415) and matrisome genes (*n* = 964) within phases and after pooling (computing union of) results for each phase. Includes counts of upregulated and downregulated genes for each phase and among all phases.

| Phase | Total DE | % DE | Upregulated | Downregulated |
| --- | --- | --- | --- | --- |
| **All genes (n = 21,407)** |  |  |  |  |
| Proliferative | 6841 | 32% | 2585 | 4256 |
| Early secretory | 2889 | 13% | 493 | 2396 |
| Mid secretory | 1327 | 6% | 312 | 1015 |
| Union of phases | 6961 | 33% | 2603 | 4357 |
| **Matrisome genes (n = 964)** |  |  |  |  |
| Proliferative | 290 | 30% | 179 | 111 |
| Early secretory | 96 | 10% | 44 | 52 |
| Mid secretory | 49 | 5% | 29 | 20 |
| Union of phases | 296 | 31% | 181 | 115 |

Next, we performed a set union of DEGs between phases (yielding a set of all genes which were differentially expressed in one or more phases) to compare the rates of differential expression in any phase between genes overall and matrisome genes. The phase-union set of DEGs contained approximately 33% of the total set of genes, while the phase-union set of DEMGs contained approximately 31% of the total matrisome genes (**Table 4**). The phase-union DEMGs were then stratified by their respective matrisome categories, and we found that ECM glycoproteins and ECM regulator were differentially expressed at higher rates than the full set of genes. In contrast, ECM-affiliated proteins were differentially expressed at the same rate as the full set of genes, and proteoglycans, secreted factors, and collagens were differentially expressed at lower rates (**Figure 3C, Table 5**). A total of 6961 unique DEGs were contained in the phase-union DEG list while 296 unique DEMGs were contained in the phase-union DEMG list (**Table 5**).

**Table 5.** Differentially expressed matrisome genes by matrisome category. Number and percentage of differentially expressed (DE) matrisome genes in each matrisome category, after performing a set union between all phases), along with number of genes in each matrisome category present in the dataset.

| Matrisome Category | Number DE | Dataset count | % DE in matrisome category |
| --- | --- | --- | --- |
| Collagens | 10 | 44 | 23% |
| ECM glycoproteins | 65 | 179 | 36% |
| ECM regulators | 84 | 230 | 37% |
| ECM-affiliated proteins | 50 | 151 | 33% |
| Proteoglycans | 10 | 35 | 29% |
| Secreted factors | 77 | 325 | 24% |

It was observed that the DEGs and DEMGs in the early and mid-secretory phases were almost entirely subsets of those found to be differentially expressed in the proliferative phase (**Figure 2A, B**). In addition, all overlapping DEGs shared by each phase were differentially expressed in the same direction, except for *ATP12A*, which was upregulated in proliferative stage samples but downregulated in mid-secretory samples (**Table 3**). Due to the fact that *ATP12A* is not defined as a matrisome gene, no DEMGs which were shared by each phase had disagreement in differential expression direction. Furthermore, DEGs and DEMGs in the mid-secretory phase seemed to be almost entirely a subset of the DEGs and DEMGs in the early-secretory phase. This indicates that the maximum dysregulation between endometriosis and normal uterine tissue occurs in the proliferative phase. It also implies that the dysregulation which occurs in the early- and mid-secretory phases is a reduced form of the dysregulation which occurs in the proliferative phase. After making this observation, we defined our final set of DEGs to be this union list, which contained genes that were differentially expressed in at least one phase of the menstrual cycle in tissue from patients with endometriosis compared to those without endometriosis across all menstrual phases. This approach identified the genes that were overall dysregulated in endometriosis regardless of menstrual phase. This final list was then used to define DEGs and DEMGs, rather than the phase-specific results.

To further explore the importance of the matrisome in tissue dysregulation between endometriosis and normal endometrium, we performed functional enrichment analysis on our final phase-union set of DEGs, defined as those genes that were differentially expressed in one or more menstrual phases. We identified several gene ontology (GO) terms (groups of functionally related genes found to be overrepresented among gene sets of interest using enrichment analysis) as enriched among our DEG list (**Figure 3D**).^10^ Enriched GO terms included functions such as focal adhesion, cell response to extracellular stimulus, and functions related to collagen and other structural ECM composition. These results further supported our decision to narrow our investigation to the matrisome gene expression specifically, instead of global gene expression.

### Machine learning classification between endometriosis and normal tissue, using matrisome gene expression

To further demonstrate that matrisome gene expression captures extensive information about endometriosis, we optimized elastic net penalized logistic regression models ^25^ to stratify diseased and normal tissue samples within each phase using only matrisome gene expression. These models achieved perfect performance in each phase (**Table 6**) measured using 5-fold cross-validated balanced accuracy.^26^ These results reinforced that gene expression within the matrisome is significantly altered in endometriosis and has substantial utility for stratifying endometriosis and normal tissue.

**Table 6.** Classifier performance. 5-fold cross validated balanced accuracy scores for classifiers within each phase which were trained to stratify endometriosis versus normal tissue. A balanced accuracy of 1 means no classification errors were made.

| Cohort | n | n<sub>endometriosis</sub> | n<sub>normal</sub> | Balanced accuracy |
| --- | --- | --- | --- | --- |
| Proliferative | 63 | 35 | 28 | 1 |
| Early secretory | 33 | 24 | 9 | 1 |
| Mid secretory | 57 | 37 | 20 | 1 |

### Stage significance analysis

To explore the dynamics of how the matrisome changes with increasing endometriosis stage, we performed several univariable and multivariable analyses, as well as weighted gene correlation network analysis (WGCNA) on the matrisome genes present in our dataset. Matrisome genes found to be significant via any of these analyses were cross-referenced with the DEMGs and classified using the following terms: DEMGs that were found to be significant via univariable or multivariable analyses were termed *stage model significant*; DEMGs that were significant via WGCNA were deemed *stage network significant*; DEMGs that were both stage model significant and stage network significant were deemed *stage significant*.

### Univariable and multivariable analysis to assess association with endometriosis stage

To investigate the relationship between individual matrisome genes and endometriosis stage, we performed the following univariable analyses: gene-wise point-biserial correlation tests between each matrisome gene and endometriosis stage ^32^ and differential gene expression (DGE) analysis among endometriosis samples comparing moderate/severe endometriosis to mild endometriosis. For multivariable analysis, we performed L^1^ penalized logistic regression, classifying endometriosis samples as moderate/severe or mild.^34, 41^ Within each phase, the L^1^ penalized logistic regression models settled on a similar number of DEMGs, and all models performed reasonably well compared to baseline values (**Table 7**). For point-biserial correlation and stage-wise DGE analysis, significance was determined based on *q*-values. For L^1^ penalized logistic regression, significance was determined based on non-zero coefficient values. These analyses yielded 214, 3, and 152 unique model significant DEMGs among the proliferative, early secretory, and mid secretory samples, respectively (**Table 8**). Of the 237 unique DEMGs identified as stage model significant within at least one phase, only 23 were not present among proliferative phase samples (**Figure 4**) and only one gene, *ANXA4*, had conflicting effects between groups. *ANXA4* was shown to be upregulated in moderate/severe versus mild endometriosis in both proliferative and mid-secretory samples but downregulated in early-secretory samples (**Table 3**). All other overlaps among model significant DEMGs between phases agreed in terms of gene effect (point-biserial correlation sign, fold-change sign in moderate/severe versus mild DGE analysis, or coefficient sign in penalized logistic regression model).

**Figure 4.**
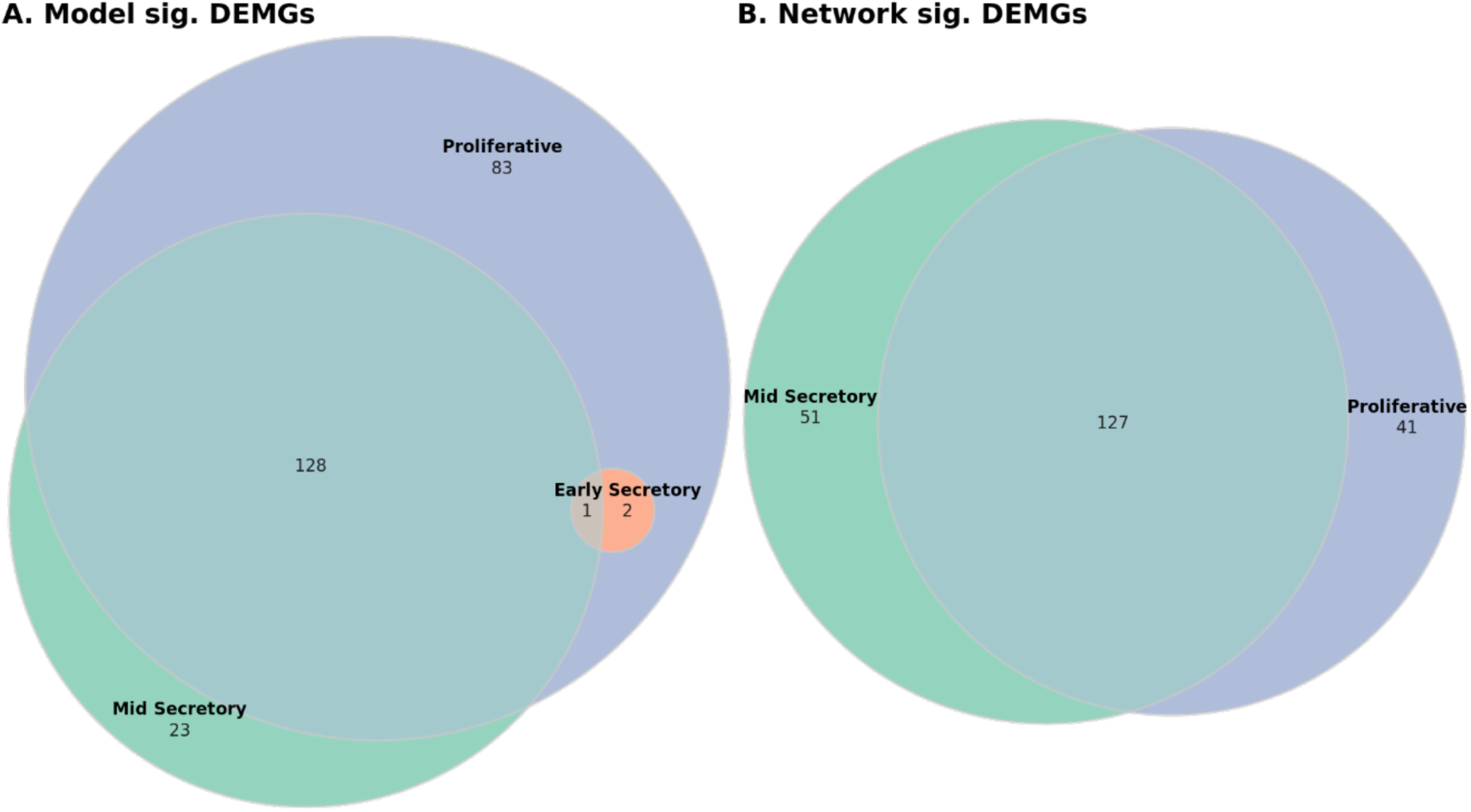
Overlaps among phases with respect to stage model significant DEMGs and stage network significant DEMGs. Overlaps among proliferative, early secretory, and mid secretory samples in terms of (**A**) stage model significant and (**B**) stage network significant differentially expressed matrisome genes (DEMGs).

**Table 7.**
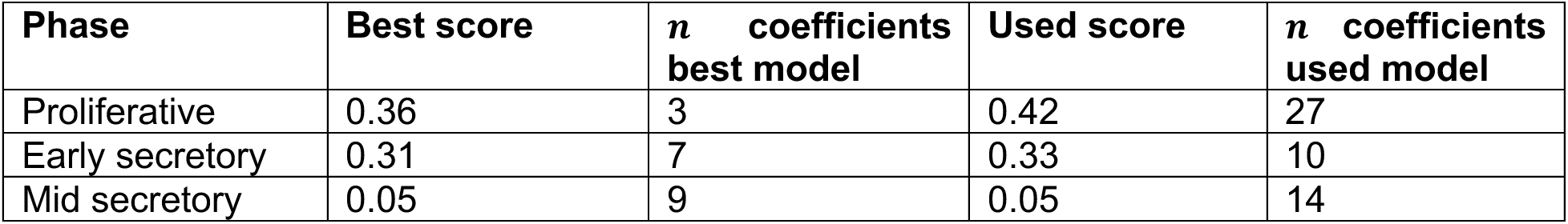
Phase-wise comparison of L1 penalized logistic regression models for classifying endometriosis stage. Performance (measured with cross-validated balanced accuracy) of the L^1^ penalized logistic regression models shows that all models outperformed the baseline of 0.5. The mid secretory models performed best. The used models were those models which had the least penalization which performed within 1 standard error of the best performance. The number of coefficients in a given model includes matrisome genes not yet filtered based on DEMG status.

**Table 8.**
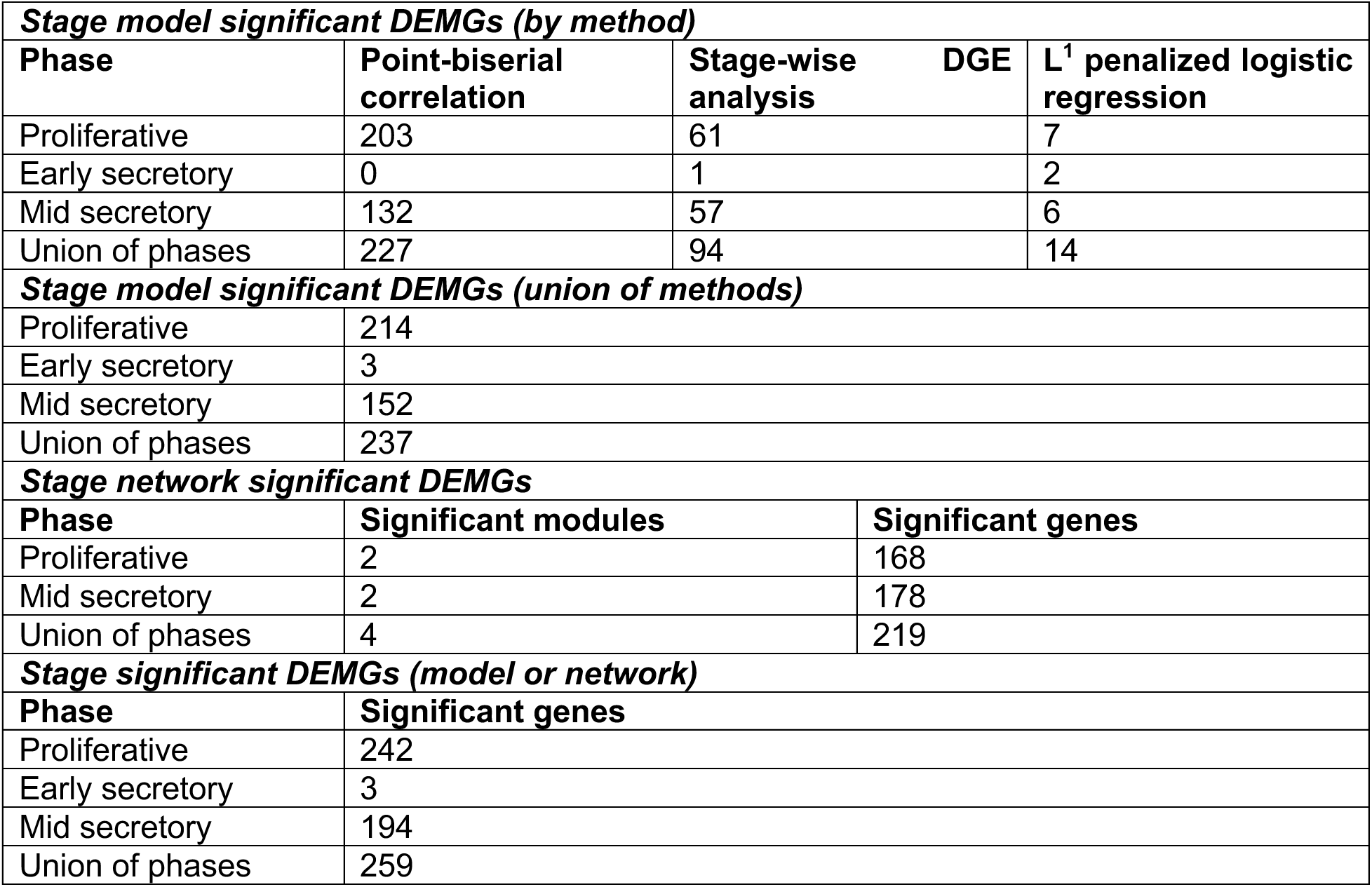
Stage model, stage network, and stage significant differentially expressed matrisome genes (DEMGs). Number of DEMGs found to be significant with respect to endometriosis stage among models, networks, or both. Results presented within phase and as union of phases. Union of methods is defined as unique genes when method results were pooled (point-biserial correlation, stage-wise differential gene expression (DGE) analysis and penalized logistic regression).

### Weighted gene correlation network analysis

WGCNA analysis identified two significant matrisome gene modules in the proliferative and two significant gene modules in the mid-secretory phases, indicating the presence of co-expressed clusters of matrisome genes within each of these two phases (**Figure 5**). Between these four modules, we identified 219 unique network significant DEMGs, with 168 and 178 network significant DEMGs found in the proliferative and mid-secretory phases, respectively (**Table 6**). As with all other analyses, extensive overlap was observed between network significant genes in the proliferative and mid-secretory phases. Gene networks were unsigned, so unlike the univariable and multivariable analyses, agreement in terms of effect direction (e.g., up or downregulation) was not assessed. Early secretory phase samples were explored with WGCNA, but no significant modules were identified, and a reasonable soft threshold value (used in WGCNA to construct topological overlap measure matrix) was not achievable for these samples.^42^ Finally, networks within phases were explored to identify hub genes, defined as genes with high levels of connectivity within their respective modules (**Figure 5**).

**Figure 5.**
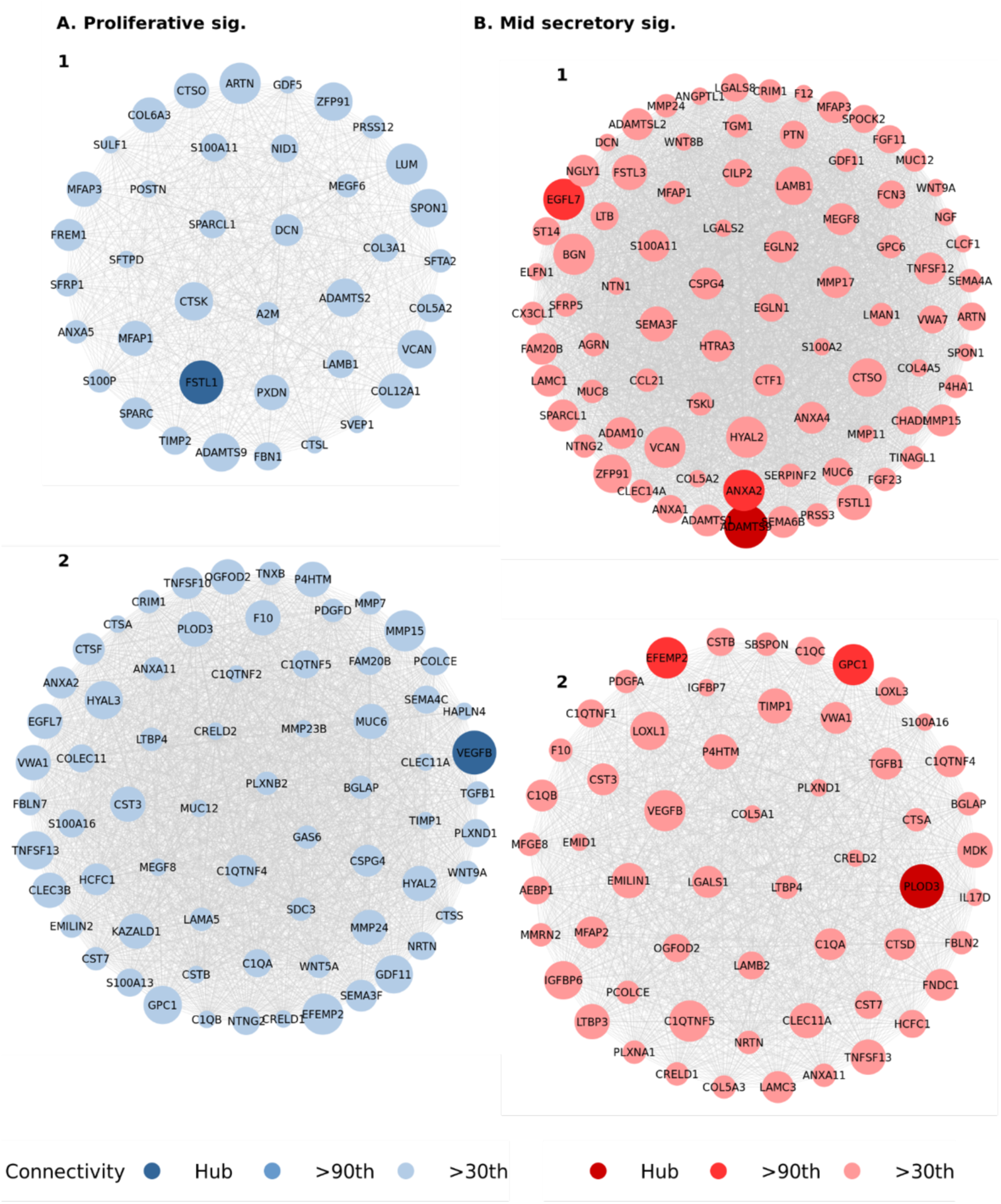
Matrisome gene network modules. Differentially expressed matrisome genes (DEMGs) whose modules, found using weighted gene correlation network analysis (WGCNA), were significantly correlated with endometriosis stage in the **(A)** proliferative and **(B)** mid secretory phases. Module genes were filtered, scaled, and shaded based on connectivity as compared to the connectivity of their module’s hub gene, the most connected DEMG in the module. Module DEMGs which were below the 30^th^ percentile in terms of connectivity are not pictured, but were utilized in our analyses. Module DEMGs which were in the 90^th^ percentile of connectivity are shaded darker than those below the 90^th^ percentile. Hub genes are shaded darkest. Connectivity was determined based on row-wise (gene-wise) sum of a given module’s adjacency matrix. Connectivity is relative to each module within each cohort, so node sizes cannot be compared between modules within the same or different cohorts.

### Stage significant genes

Both the model and network analyses evaluated gene significance with respect to disease stage. Thus, results for these analyses were combined within each menstrual phase. As with our DGE analysis between diseased and normal endometrium, the majority of stage significant DEMGs across all menstrual phases were a subset of those that were significant within the proliferative phase. Therefore, the DEMGs that were both network and model significant were pooled between phases. Early-secretory had very few stage significant DEMGs, which may be due to reduced statistical power, as there were fewer samples in the early-secretory phase compared to the other menstrual phases (**Table 9**). Alternatively, the smaller number of stage significant DEMGs could be due to an underlying biological mechanism.

**Table 9.** Sample counts within each phase stratified by endometriosis status. Number of samples within each of the three menstrual cycle phases in our dataset, stratified by whether the tissue sample is normal or has mild or moderate/severe endometriosis.

| Phase | Endometriosis status | *n* |
| --- | --- | --- |
| Proliferative | Normal | 28 |
|  | Mild endometriosis | 12 |
|  | Moderate/severe endometriosis | 23 |
| Early secretory | Normal | 9 |
|  | Mild endometriosis | 6 |
|  | Moderate/severe endometriosis | 18 |
| Mid secretory | Normal | 20 |
|  | Mild endometriosis | 9 |
|  | Moderate/severe endometriosis | 28 |

Among the stage significant DEMGs were 60 secreted factors including chemokines *CCL3*, *CCL5*, *CCL14*, *CCL21*, *CX3CL1*, and *CXCL14*, interleukins *IL13, IL15*, and *IL17C*, growth factors *NGF*, *PDGFA*, *TGFB1*, *TNF*, and *VEGFB*, 65 ECM regulators including ADAM metallopeptidase, matrix metallopeptidase, cathepsin, and lysyl oxidase families, 56 glycoproteins including agrin, elastin, fibrillin, laminin, and matrillin families, 41 ECM affiliated proteins including lectin and mucin families, 10 genes related to collagen including the COL4A and COL5A families, and 8 proteoglycans including decorin, podocin, and versican (**Figure 6, Supplemental Table 1**).

**Figure 6.**
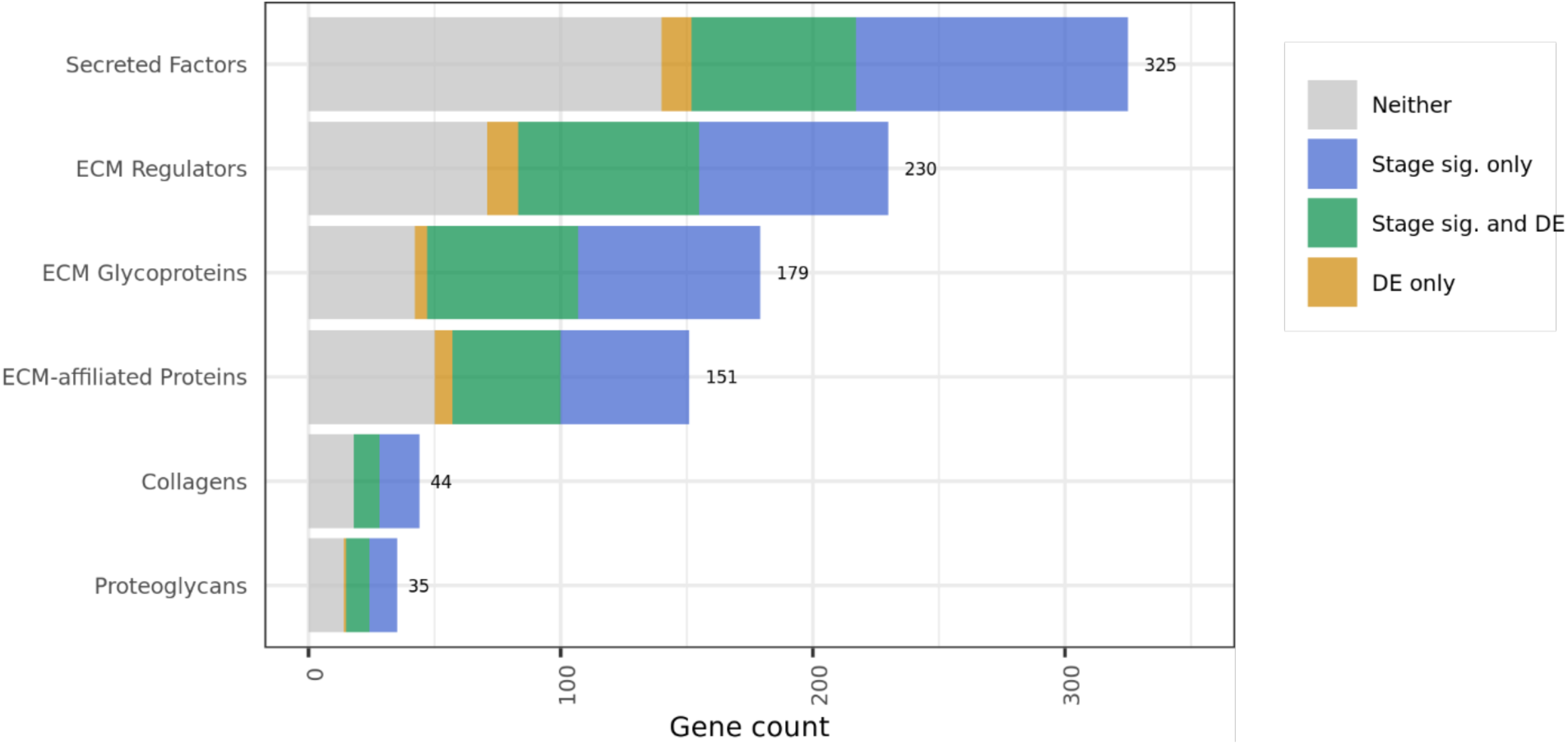
Matrisome category differential expression and stage significance breakdown. Visualization of the overlap between matrisome genes which are differentially expressed (DE) in endometriosis versus normal tissue and matrisome genes which have inferential significance with respect to endometriosis stage. The green area (overlap between DE and stage significant matrisome genes) represents the stage significant differentially expressed matrisome genes which were the subject of a large portion of our analysis.

### Functional enrichment and pathway analysis

As the majority of DEMGs were stage significant, the enriched GO terms and pathways among DEMGs were largely preserved among stage significant DEMGs. Among DEMGs and stage significant DEMGs, GO terms such as extracellular matrix, extracellular structure, and external encapsulating structure organization were highly enriched due to dysregulation of ADAM and ADAMTS family genes, collagens, laminins, matrix metallopeptidases, and others (**Figure 7A**). Basement membrane, cytokine activity, growth factor activity, and glycosaminoglycan binding were also significantly enriched among DEMGs and stage significant DEMGs (**Figure 7A**).

**Figure 7.**
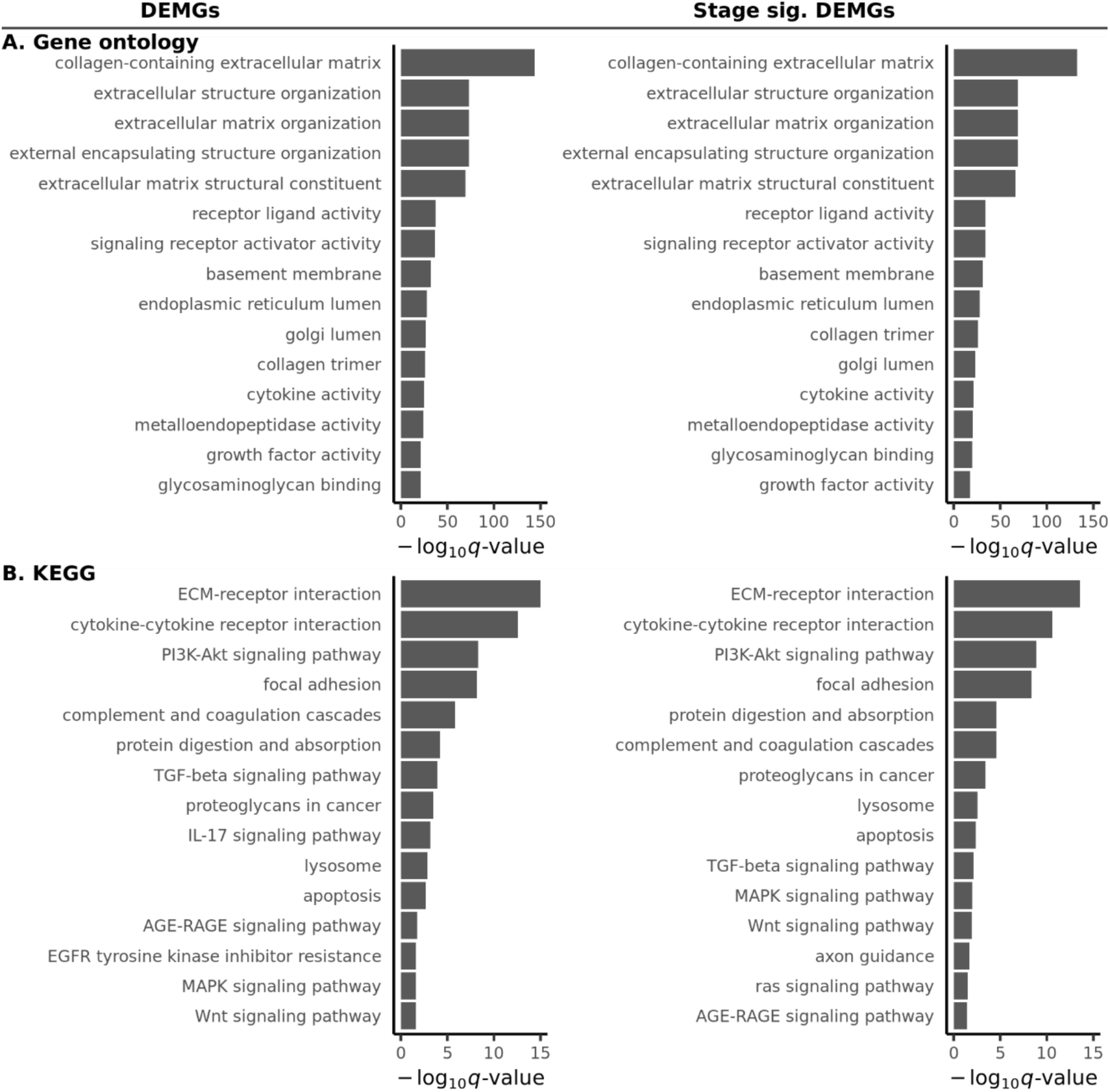
Functional enrichment (Gene ontology) and pathway (KEGG) analysis results. Top 15 most significantly enriched **(A)** gene ontology terms and **(B)** KEGG pathways among differentially expressed matrisome genes (DEMGs) and stage significant DEMGs. Value of − *log*_1;_(*q*) > 1.3 indicates significance (*q* < 0.05).

The Kyoto Encyclopedia of Genes and Genomes (KEGG) is a knowledge base which connects established genomic information with high order functional behavior among genes, defining pathways which describe critical cellular processes ^11^. Pathways that were significantly enriched among DEMGs and stage significant DEMGs included ECM-receptor interaction, cytokine-cytokine receptor interaction, PI3K-Akt signaling, focal adhesion, complement and coagulation cascades, protein digestion and absorption, TGF-β signaling, lysosome, AGE-RAGE signaling, MAPK signaling, Wnt signaling, and axon guidance (**Figure 7B**). Only the IL-17 signaling, EGFR tyrosine kinase inhibitor resistance, HIF-1 signaling, and glycosaminoglycan degradation pathways were enriched among DEMGs only, and not stage significant DEMGs.

## Discussion

In summary, we analyzed the relationship between matrisome gene expression and the presence and stage of endometriosis. First, we identified genes that were differentially expressed between endometriosis and normal tissue and established that ECM-related GO terms were significantly enriched among these genes. Next, we demonstrated that machine learning models could distinguish between normal and endometriosis tissue using matrisome gene expression data alone. We then identified matrisome genes and gene networks that had inferential significance for the progression of endometriosis and used these results to identify dysregulated pathways and gene ontology terms.

While the endometrium is a highly dynamic tissue and samples from different menstrual cycle phases should likely be analyzed separately, our results suggest that most dysregulation in the early- and mid-secretory phases can be viewed as subsets of the dysregulation in the proliferative phase. We observed that proliferative phase samples had the most differentially expressed genes overall and the most differentially expressed matrisome genes. We also observed that proliferative phase samples yielded the most extensive sets of dysregulated matrisome genes when analyzing the relationship between matrisome gene expression and mild versus moderate endometriosis. This could mean that the proliferative phase of the menstrual cycle is the ideal time for performing protein or gene expression-based analysis on tissue from people with endometriosis. Furthermore, we found that dysregulation of matrisome genes in endometrial tissue from people with endometriosis compared to those without may be a good indicator of significance for inference about the progression of endometriosis. This finding implies that the expression of the same matrisome genes can be used to study both the onset and progression of endometriosis.

Our work confirmed and consolidated previous findings on the dysregulation of matrisome genes and pathways in endometriosis. For example, similar to previous bioinformatics studies of endometriosis, we found that ECM-receptor interactions, cytokine-cytokine receptor interactions, immune-stromal cell interactions, coagulation cascades, and TGF-β signaling were dysregulated in endometriosis tissue compared to healthy endometrium.^7, 43–46^ We also found that inflammatory and neurotransmission cytokines and pathways were correlatively dysregulated, which is in line with studies that have investigated neuroinflammation in endometriosis patients.^47, 48^ The PI3K-Akt signaling pathway was significantly enriched in endometriosis samples and inferentially significant for endometriosis stage. Upregulation of PI3K-Akt has been reported in animal models of endometriosis as well as eutopic endometrium samples from people with endometriosis.^49, 50^ The AGE-RAGE was also dysregulated and significant for endometriosis stage, which has been linked to endometriosis pathogenesis as well as oxidative stress, inflammation, apoptosis, and angiogenesis.^51^ Lastly, our work confirmed that both the MAPK and Wnt signaling pathways are important for the onset and progression of endometriosis, which have been implicated in endometriosis pathology through *in vitro* experiments.^52, 53^

While combining results from different phases for enrichment analysis was justifiable given the extensive inter-phase overlaps observed, this may obfuscate more granular characteristics of each phase. Additionally, our analyses were limited to only eutopic samples of normal and endometriosis endometrium. This constraint allowed us to control for variation attributable to the tissue of origin but prevented us from considering matrisome characteristics of ectopic endometrium. As endometriosis datasets grow in size and tissue diversity, matrisome expression analysis of ectopic endometrium could be an area of interest for future work. Future work could also attempt to deconvolve the activity of specific cell types involved in the matrisome dysregulation we observed, similar to the use of CIBERSORT and xCell in the work by Poli-Neto.^4, 54, 55^ Finally, we only investigated matrisome genes that were dysregulated between normal and endometriosis tissue overall when assessing significance for endometriosis progression inference. Future work could expand on our unfiltered analysis and explore results for matrisome genes which were significant for endometriosis progression without cross-referencing for differential expression in disease overall.

This work builds upon our previous work that used a similar approach to analyze the relationships between matrisome gene expression and gynecological cancers.^56^ This analysis pipeline represents a clear and consolidated application of many of the most influential and well-established methods for analyzing transcriptional data and machine learning methods to determine which genes and gene networks are significant for the onset and progression of disease. This provides individuals with expertise in ECM biology or tissue engineering but little expertise in computer science with an overview of how to analyze their datasets and identify matrisome components of interest for their applications.

Overall, the work presented here is one of the most comprehensive omics analyses of endometriosis data currently available, and to our knowledge, the only such study which focuses on exploring matrisome dysregulation of endometriosis. Our results reinforce and expand upon previous findings related to gene expression dysregulation in endometriosis and hold significant value for future drug discovery and tissue engineering research focused on endometriosis.

## Conflicts of interest

The authors have no conflicts to declare.

## IRB statement

IRB approval was not required for this study, as it was a meta-analysis of existing publicly available data.

## Data and code availability

All data and code are available on the Fogg Lab Github (https://github.com/fogg-lab/)

## Notes

### Competing Interest Statement

The authors have declared no competing interest.

## References

1. Parasar P, Ozcan P, Terry KL. Endometriosis: Epidemiology, Diagnosis and Clinical Management. Curr Obstet Gynecol Rep. 2017 Mar;6(1):34–41. PMCID: PMC5737931

2. Hansen KA, Eyster KM. Genetics and genomics of endometriosis. Clin Obstet Gynecol. 2010 Jun;53(2):403–412. PMCID: PMC4346178

3. Daftary GS, Zheng Y, Tabbaa ZM, Schoolmeester JK, Gada RP, Grzenda AL, et al. A Novel Role of the Sp/KLF Transcription Factor KLF11 in Arresting Progression of Endometriosis. PLOS ONE. Public Library of Science; 2013 Mar 28;8(3):e60165.

4. Poli-Neto OB, Meola J, Rosa-E-Silva JC, Tiezzi D. Transcriptome meta-analysis reveals differences of immune profile between eutopic endometrium from stage I-II and III-IV endometriosis independently of hormonal milieu. Sci Rep. 2020 Jan 15;10(1):313. PMCID: PMC6962450

5. Barnhart K, Dunsmoor-Su R, Coutifaris C. Effect of endometriosis on in vitro fertilization. Fertil Steril. 2002 Jun;77(6):1148–1155. PMID: 12057720

6. Bałkowiec M, Maksym RB, Włodarski PK. The bimodal role of matrix metalloproteinases and their inhibitors in etiology and pathogenesis of endometriosis (Review). Mol Med Rep. 2018 Sep;18(3):3123–3136. PMCID: PMC6102659

7. Yu L, Shen H, Ren X, Wang A, Zhu S, Zheng Y, et al. Multi-omics analysis reveals the interaction between the complement system and the coagulation cascade in the development of endometriosis. Sci Rep. 2021 Jun 7;11:11926. PMCID: PMC8185094

8. Bonnans C, Chou J, Werb Z. Remodelling the extracellular matrix in development and disease. Nat Rev Mol Cell Biol. Nature Publishing Group; 2014 Dec;15(12):786–801.

9. Naba A, Clauser KR, Hoersch S, Liu H, Carr SA, Hynes RO. The matrisome: in silico definition and in vivo characterization by proteomics of normal and tumor extracellular matrices. Mol Cell Proteomics. 2012 Apr;11(4):M111.014647. PMCID: PMC3322572

10. Ashburner M, Ball CA, Blake JA, Botstein D, Butler H, Cherry JM, et al. Gene Ontology: tool for the unification of biology. Nat Genet. Nature Publishing Group; 2000 May;25(1):25–29.

11. Kanehisa M, Goto S. KEGG: kyoto encyclopedia of genes and genomes. Nucleic Acids Res. 2000 Jan 1;28(1):27–30. PMCID: PMC102409

12. R: The R Project for Statistical Computing [Internet]. [cited 2023 Mar 2]. Available from: https://www.r-project.org/

13. Talbi S, Hamilton AE, Vo KC, Tulac S, Overgaard MT, Dosiou C, et al. Molecular Phenotyping of Human Endometrium Distinguishes Menstrual Cycle Phases and Underlying Biological Processes in Normo-Ovulatory Women. Endocrinology. 2006 Mar 1;147(3):1097– 1121.

14. Burney RO, Talbi S, Hamilton AE, Vo KC, Nyegaard M, Nezhat CR, et al. Gene expression analysis of endometrium reveals progesterone resistance and candidate susceptibility genes in women with endometriosis. Endocrinology. 2007 Aug;148(8):3814–3826. PMID: 17510236

15. Hever A, Roth RB, Hevezi P, Marin ME, Acosta JA, Acosta H, et al. Human endometriosis is associated with plasma cells and overexpression of B lymphocyte stimulator. Proc Natl Acad Sci U S A. 2007 Jul 24;104(30):12451–12456. PMCID: PMC1941489

16. Tamaresis JS, Irwin JC, Goldfien GA, Rabban JT, Burney RO, Nezhat C, et al. Molecular classification of endometriosis and disease stage using high-dimensional genomic data. Endocrinology. 2014 Dec;155(12):4986–4999. PMCID: PMC4239429

17. Irizarry RA, Bolstad BM, Collin F, Cope LM, Hobbs B, Speed TP. Summaries of Affymetrix GeneChip probe level data. Nucleic Acids Res. 2003 Feb 15;31(4):e15. PMCID: PMC150247

18. Irizarry RA, Hobbs B, Collin F, Beazer-Barclay YD, Antonellis KJ, Scherf U, et al. Exploration, normalization, and summaries of high density oligonucleotide array probe level data. Biostatistics. 2003 Apr;4(2):249–264. PMID: 12925520

19. Gautier L, Cope L, Bolstad BM, Irizarry RA. affy—analysis of Affymetrix GeneChip data at the probe level. Bioinformatics. 2004 Feb 12;20(3):307–315.

20. Leek JT. svaseq: removing batch effects and other unwanted noise from sequencing data. Nucleic Acids Research. 2014 Dec 1;42(21):e161–e161.

21. Li J, Bushel PR, Chu TM, Wolfinger RD. Principal Variance Components Analysis: Estimating Batch Effects in Microarray Gene Expression Data. Batch Effects and Noise in Microarray Experiments [Internet]. John Wiley & Sons, Ltd; 2009 [cited 2023 Mar 2]. p. 141–154. Available from: https://onlinelibrary.wiley.com/doi/abs/10.1002/9780470685983.ch12

22. niehs.nih.gov> PB <bushel at. pvca: Principal Variance Component Analysis (PVCA) [Internet]. Bioconductor version: Release (3.16); 2023 [cited 2023 Mar 2]. Available from: https://bioconductor.org/packages/pvca/

23. Davis S, Meltzer PS. GEOquery: a bridge between the Gene Expression Omnibus (GEO) and BioConductor. Bioinformatics. 2007 Jul 15;23(14):1846–1847.

24. Hynes RO, Naba A. Overview of the matrisome--an inventory of extracellular matrix constituents and functions. Cold Spring Harb Perspect Biol. 2012 Jan 1;4(1):a004903. PMCID: PMC3249625

25. Regularization and variable selection via the elastic net - Zou - 2005 - Journal of the Royal Statistical Society: Series B (Statistical Methodology) - Wiley Online Library [Internet]. [cited 2023 Mar 2]. Available from: https://rss.onlinelibrary.wiley.com/doi/full/10.1111/j.1467-9868.2005.00503.x

26. The Balanced Accuracy and Its Posterior Distribution | IEEE Conference Publication | IEEE Xplore [Internet]. [cited 2023 Mar 2]. Available from: https://ieeexplore.ieee.org/document/5597285

27. Pedregosa F, Varoquaux G, Gramfort A, Michel V, Thirion B, Grisel O, et al. Scikit-learn: Machine Learning in Python. Journal of Machine Learning Research. 2011;12(85):2825– 2830.

28. Hutter F, Hoos HH, Leyton-Brown K. Sequential Model-Based Optimization for General Algorithm Configuration. In: Coello CAC, editor. Learning and Intelligent Optimization. Berlin, Heidelberg: Springer; 2011. p. 507–523.

29. Head T, MechCoder, Louppe G, Shcherbatyi I, fcharras, Vinícius Z, et al. scikit-optimize/scikit-optimize: v0.5.2 [Internet]. Zenodo; 2018 [cited 2023 Mar 2]. Available from: https://zenodo.org/record/1207017/export/xd

30. Ritchie ME, Phipson B, Wu D, Hu Y, Law CW, Shi W, et al. limma powers differential expression analyses for RNA-sequencing and microarray studies. Nucleic Acids Res. 2015 Apr 20;43(7):e47. PMCID: PMC4402510

31. Storey JD. The positive false discovery rate: a Bayesian interpretation and the q-value. The Annals of Statistics. Institute of Mathematical Statistics; 2003 Dec;31(6):2013–2035.

32. Kornbrot D. Point Biserial Correlation. Encyclopedia of Statistics in Behavioral Science [Internet]. John Wiley & Sons, Ltd; 2005 [cited 2023 Mar 2]. Available from: https://onlinelibrary.wiley.com/doi/abs/10.1002/0470013192.bsa485

33. Langfelder P, Horvath S. WGCNA: an R package for weighted correlation network analysis. BMC Bioinformatics. 2008 Dec 29;9(1):559.

34. Friedman J, Hastie T, Tibshirani R, Narasimhan B, Tay K, Simon N, et al. glmnet: Lasso and Elastic-Net Regularized Generalized Linear Models [Internet]. 2022 [cited 2023 Mar 2]. Available from: https://CRAN.R-project.org/package=glmnet

35. Zhang B, Horvath S. A general framework for weighted gene co-expression network analysis. Stat Appl Genet Mol Biol. 2005;4:Article17. PMID: 16646834

36. Langfelder P, Mednet Sh. Tutorials for the WGCNA package [Internet]. Tutorials for the WGCNA package. 2011 [cited 2023 Mar 2]. Available from: https://horvath.genetics.ucla.edu/html/CoexpressionNetwork/Rpackages/WGCNA/Tutorials/

37. Wu T, Hu E, Xu S, Chen M, Guo P, Dai Z, et al. clusterProfiler 4.0: A universal enrichment tool for interpreting omics data. Innovation (Camb). 2021 Aug 28;2(3):100141. PMCID: PMC8454663

38. Gene Ontology (GO) database and informatics resource | Nucleic Acids Research | Oxford Academic [Internet]. [cited 2023 Mar 2]. Available from: https://academic.oup.com/nar/article/32/suppl_1/D258/2505186

39. gmail.com> JTL <jtleek at, bu.edu> WEJ <wej at, jhsph.edu> HSP <hiparker at, jhmi.edu> EJF <ejfertig at, jhsph.edu> AEJ <ajaffe at, gmail.com> YZ <zhangyuqing pkusms at, et al. sva: Surrogate Variable Analysis [Internet]. Bioconductor version: Release (3.16); 2023 [cited 2023 Mar 2]. Available from: https://bioconductor.org/packages/sva/

40. Wang W, Vilella F, Alama P, Moreno I, Mignardi M, Isakova A, et al. Single-cell transcriptomic atlas of the human endometrium during the menstrual cycle. Nat Med. Nature Publishing Group; 2020 Oct;26(10):1644–1653.

41. Sparse Multinomial Logistic Regression via Bayesian L1 Regularisation. 2007 Sep 7 [cited 2023 Mar 2]; Available from: https://direct.mit.edu/books/book/3168/chapter/87394/Sparse-Multinomial-Logistic-Regression-via

42. WGCNA package: Frequently Asked Questions [Internet]. [cited 2023 Mar 2]. Available from: https://horvath.genetics.ucla.edu/html/CoexpressionNetwork/Rpackages/WGCNA/faq.html

43. Differentially expressed genes in human endometrial endothelial cells derived from eutopic endometrium of patients with endometriosis compared with those from patients without endometriosis | Human Reproduction | Oxford Academic [Internet]. [cited 2023 Mar 2]. Available from: https://academic.oup.com/humrep/article/22/12/3159/2384929

44. Liu F, Lv X, Yu H, Xu P, Ma R, Zou K. In search of key genes associated with endometriosis using bioinformatics approach. Eur J Obstet Gynecol Reprod Biol. 2015 Nov;194:119–124. PMID: 26366788

45. Ping S, Ma C, Liu P, Yang L, Yang X, Wu Q, et al. Molecular mechanisms underlying endometriosis pathogenesis revealed by bioinformatics analysis of microarray data. Arch Gynecol Obstet. 2016 Apr;293(4):797–804. PMID: 26354330

46. Symons LK, Miller JE, Kay VR, Marks RM, Liblik K, Koti M, et al. The Immunopathophysiology of Endometriosis. Trends Mol Med. 2018 Sep;24(9):748–762. PMID: 30054239

47. Arellano Estrada C, Barcena de Arellano ML, Schneider A, Mechsner S. Neuroimmunomodulation in the pathogenesis of endometriosis. Brain, Behavior, and Immunity. 2013 Feb 15;29:S2.

48. Wei Y, Liang Y, Lin H, Dai Y, Yao S. Autonomic nervous system and inflammation interaction in endometriosis-associated pain. Journal of Neuroinflammation. 2020 Mar 7;17(1):80.

49. Mu L, Zheng W, Wang L, Chen XJ, Zhang X, Yang JH. Alteration of focal adhesion kinase expression in eutopic endometrium of women with endometriosis. Fertil Steril. 2008 Mar;89(3):529–537. PMID: 17543958

50. Fibrinogen alpha chain promotes the migration and invasion of human endometrial stromal cells in endometriosis through focal adhesion kinase/protein kinase B/matrix metallopeptidase 2 pathway† | Biology of Reproduction | Oxford Academic [Internet]. [cited 2023 Mar 2]. Available from: https://academic.oup.com/biolreprod/article/103/4/779/5874328

51. Fujii EY, Nakayama M, Nakagawa A. Concentrations of receptor for advanced glycation end products, VEGF and CML in plasma, follicular fluid, and peritoneal fluid in women with and without endometriosis. Reprod Sci. 2008 Dec;15(10):1066–1074. PMID: 19088375

52. Yoshino O, Osuga Y, Hirota Y, Koga K, Hirata T, Harada M, et al. Possible pathophysiological roles of mitogen-activated protein kinases (MAPKs) in endometriosis. Am J Reprod Immunol. 2004 Nov;52(5):306–311. PMID: 15550066

53. Matsuzaki S, Darcha C. Involvement of the Wnt/β-catenin signaling pathway in the cellular and molecular mechanisms of fibrosis in endometriosis. PLoS One. 2013;8(10):e76808. PMCID: PMC3790725

54. Aran D, Hu Z, Butte AJ. xCell: digitally portraying the tissue cellular heterogeneity landscape. Genome Biology. 2017 Nov 15;18(1):220.

55. Newman AM, Steen CB, Liu CL, Gentles AJ, Chaudhuri AA, Scherer F, et al. Determining cell type abundance and expression from bulk tissues with digital cytometry. Nat Biotechnol. 2019 Jul;37(7):773–782. PMCID: PMC6610714

56. Cook CJ, Miller AE, Barker TH, Di Y, Fogg KC. Characterizing the extracellular matrix transcriptome of cervical, endometrial, and uterine cancers. Matrix Biology Plus. 2022 Jul 16;100117.

